# FirstSV: Fast and Accurate Approach of Structural Variations Detection for Short DNA fragments

**DOI:** 10.1101/415059

**Authors:** Jia Shen, Qiyang Zuo, Rongliang Wang, Xiang Li, Yuanhua Tang

**Author notes:** these authors contributed equally to this work (Jia Shen, Qiyang Zuo).

## Abstract

Structural variations caused by gene fusion represent a major class of somatically acquired variations in human malignancies, and include deletions, inversions, and translocations. Short fragmented reads are the main source of data from 2nd-generation sequencing, and detecting structural variations from this type of data is different from that of 1st-generation sequencing, where the read length is much longer. Current detection methods are low in specificity and are inefficient. We developed a hybrid algorithm, FirstSV, to meet the clinical demand for fast and accurate structural variation detection. Its main features include cluster analysis, realignment, and local assembly. FirstSV was validated with simulated data, with data from real patient samples, with data from standard testing samples, and with downloaded public data sets. FirstSV outperforms public-available methods in terms of sensitivity, precision, and operational efficiency. FirstSV is freely available at https://github.com/shenjia1/FirstSV.

## Introduction

Structural variations (SVs) are an important type of genetic polymorphisms, including large deletions, inversions, and translocations. SVs resulting from gene fusions represent a major class of somatically acquired structural variation in human malignancies, for example, the ALK gene fusions occurred quite frequently in non-small-cell lung carcinoma (NSCLC)^1,2,3^. Detecting SVs is different from detecting single nucleotide polymorphisms, or short indels. So far, researchers have developed several algorithms for such a task.

Current detection methods based on next-generation sequencing reads can be classified into the following categories: Clustering, split-reads alignment, contig assembly, and hybrid approaches^4,5^. Clustering methods transform the structural variations detections to grouping reads that support the same structural variations. It can detect SVs quickly but can not find the exact breakpoints. Breakdancer, a classical clustering methods, with relatively high sensitivity but low specificity, can only handle discordant read pairs, not the unpaired reads^6^. Seeksv is a split-reads alignment algorithm. It aligns soft-clipped reads and one-end-anchored reads to genome^7^. It needs many split-clipped reads to locate breakpoints and to identify the exact breakpoints. Split-reads alignment suits current sequencing data with increasing read length and is also adopted in our study^8^. Contig assembly method first assembles anomalously mapped reads to form contigs and remapped contigs to the reference genome to detect the breakpoints. In a sense, contigs assembly methods are the most suitable algorithm for breakpoint detection as these methods can remove mapping errors and find breakpoints from short reads. However, the assembly algorithm was relatively inefficient, and an initial filtering of candidate areas was needed. Current relative mature assembly algorithms, such as Soapdenovo2^9^, SGA^10^ and CAP^11^, were mostly designed for whole genome assembly.

Local assembly task was different from whole genome assembly as the whole genome assembly was much more complex than local assembly, and the resulted assembly output were not ideal. There were also some assembly software packages designed for small fragments assembly or local assembly, such as TIGRA^12^, Fermi^13^ and novoBreak^14^. Fermi, based on the overlap layout-consensus-assembly algorithm, was adopted in this study for local assembly. Hybrid approaches usually combine two or more algorithms to identify the breakpoints. factera, a hybrid algorithm, improved the efficiency of detection with high sensitivity, clustered the discordant reads to discover potential regions of breakpoints and realigned the softclipped reads in these regions to the reference genome to confirm the extract breakpoints. Unfortunately, it was not designed to handle mapping errors^15^.

Most of the detection algorithms mentioned above were designed to detect structural variations in whole genome. However, in clinical settings, we do not need whole-genome assembly. We only need to identify breakpoints in very specific regions of the genome, as these regions are the hotspots of structural mutations. Targeted sequencing of the specific regions in the genome could significantly reduce the cost, targeted toward problematic regions. The frequency of somatic structural variations was often too low to detect, especially in cfDNA data, which required much higher sensitivity. Existing software packages were adjusted and used in clinical detections, but the overall results were not good.

For these reasons, we developed a new method by combining clustering alignment, realignment, and local assembly, and we named it FirstSV. FirstSV first cluster discordant reads and assemble sequences near the potential breakpoints, and then realign the assembled sequences to the reference genome to find the exact breakpoints. Split-reads alignment was utilized to locate the potential breakpoints if the read length was long enough to deal with. Each pair of incorrect mapping reads were analyzed with high efficiency to ensure high sensitivity. Some filtering rules were utilized to avoid most of false positives and to ensure high specificity. In order to validate FirstSV, public data sets, lung cancer sample data, and commercial standard sample data were used to test. The test results show that FirstSV was superior to other pieces of software in terms of detection rate and efficiency, and it achieved good performance in specificity and sensitivity.

## Results

### Characteristics of cfDNA samples

In clinical gene testing, cfDNA samples were different from gDNA samples. Because cfDNA fragments are only about 167bp in length. cfDNA samples were sequenced directly without fragmentation^16,17^. The 150bp, paired-end sequences were generated using Illumina NextSeq500 sequencer and the average sequencing depth per sample was 5000×. Since the sequencing read length was close to the cfDNA fragment length, pair of sequences were mostly overlapping. Because the cfDNA fragments were very short, traditional methods of analyzing discordant reads signatures could not detect structural variations. To solve the problem, the softclip reads signature was first analyzed in candidate regions, and then in the local assembly of candidate regions in FirstSV (Figure 1). A softclip signature means that a sequence is unclipped even it maps only partially to the genome.

**Figure 1.**
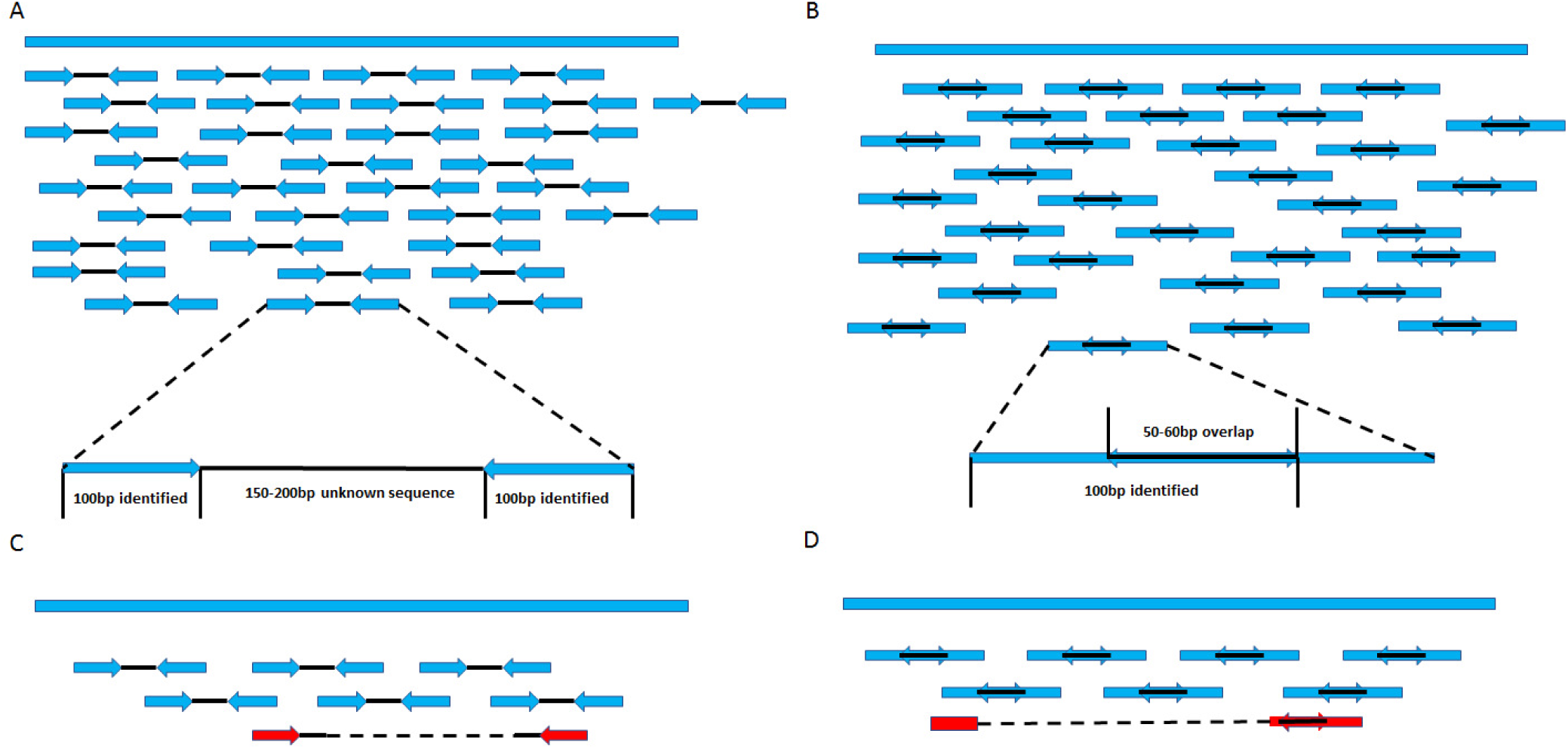
Comparison of data characteristics of cfDNA and gDNA.gDNA samples would be randomly fragmented before sequencing. The length of gDNA fragments after interrupting was about 500bp. As the second-generation sequencing read length is very limited, only the two ends of DNA fragments could be sequenced. As shown in (A), supposing that length of DNA fragment was range from 350bp to 500bp and sequencing read length is 100bp, the middle sequence of DNA fragment was not available. DNA fragment size in cfDNA samples was about 167bp and cfDNA samples could be directly sequenced and all information for the DNA fragment could be obtained (B). Most of the breakpoints were in the unknown area of gDNA fragments, so exact location of breakpoints may be missed by the sequencing data. Pair-end reads might map to discordant locations, which could help finding the structural variations. The red fragment is a fragment of structural variation. Because the gDNA fragment is too long to be sequenced through, the exact location of breakpoints may be hard to find (C). Of course, there might be occasions where the fragments contain the breakpoints within 100bp of the ends, and can be determined using local assembly. In cfDNA samples, the structural variation could be directly found by the alignment of the sequences, which was called the softclip signature. The red fragment is a fragment with structural variation. The cfDNA fragment is so short that the exact breakpoints can be determined quite often.

### Analysis of simulated data

In order to evaluate out software, simulated data were used. We used in-house software to simulate 147 known structural variations in COSMIC databases, including 34 deletions, 38 inversions, and 75 translocations. Then wgsim^20^ was used to simulate PE150 sequencing data, of which average insert size was 166bp and the standard deviation was 30. We simulated the sequencing data of different depth gradients of 10, 20, 50, 100, 500, 1000 and 5000 respectively. We also simulated the sequencing data with mutation rate at 0.001, 0.005, 0.01, 0.05, 0.2, 0.5 and 1 on 5000×. We ran the same data set using 4 distinct software pieces: FirstSV, svaba(https://github.com/walaj/svaba), brekdancer, and breakmer. The outputs from each method is shown in Table 4. One can see that our software program is superior to other software pieces in sensitivity and precision. Our software detected all the structural variations, with the least amount of false positives for all cases of SVs simulated (deletion, inversion, translocation).

### Analysis of standard samples

To test the reproducibility of FirstSV, Horizon’s Structural Multiplex Reference Standard was chosen. This standard is designed to challenge one’s bioinformatics workflow by providing validated copy number variants/amplifications, translocations, and large insertions/deletions. The Structural Multiplex includes 9 ddPCR-validated mutations, with most of them centered at 5% allelic frequency. Highlight features of the Structural Multiplex include RET and ROS1 fusion variants, MYC-N and MET focal amplifications, and a BRCA2 variant. The Structural Multiplex is also available in cfDNA (HD786) and FFPE (HD789). We used cfDNA standard sample to test FirstSV. Sequencing and data preprocessing was the same as Methods.. Two repeated sequencing of the standard were tested and both of them got the expected results, in which the detection frequency was close to the expected result (Table 1).

**Table 1.** Analysis of standard samples by FirstSV. The frequency of structural variations was contained in the standard. *True-positive represented the standard frequency of SV.None of false positive was detected and the expected structural variations were SLC34A2-ROS1 and CCDC6-RET.

| Type | First | Second | *True-positive |
| --- | --- | --- | --- |
| SLC34A2-ROS1 | 0.048 | 0.06 | 0.056 |
| CCDC6-RET | 0.05 | 0.04 | 0.05 |

### Analysis of lung cancer samples

To evaluate the performance of FirstSV in clinical settings, eight NSCLC tumor samples, consisting of four cfDNA samples and four cancer tissue samples, harbored a known rearrangement in ALK-EML4, were chosen. Tissue DNA was fragmented to a mean size of 200bp by sonication, whereas cfDNA samples was not sonicated. Details on data preprocessing followed the methods mentioned above. FirstSV had a 100% detection rate, and an average of 6.1 CPU minutes in run time were per sample. The other software pieces, factera, seeksv, svaba, breakdancer were run by their recommended default parameter settings, with corresponding detection rates at 12.5%, 37.5%, 87.5%, and 75% respectively (Table 2). FirstSV has the highest detection rate. Except for factera and breakdancer, FirstSV was also better at efficiency than other software pieces, suggesting that FirstSV was superior to other software pieces in targeted sequencing data with high depth, especially in detecting structural variations of cfDNA sample data.

**Table 2.** Structure variation detection of NSCLC tumor samples by multiple software packages. * indicates the breakpoints produced by breakdancer were not exact positions. The expect number of true SVs is 8.

| Softwares | Counts of Detection | Time(min) | Detection Rate | Additional Calls Per Sample |
| --- | --- | --- | --- | --- |
| FirstSV | 8 | 6.1 | 1 | 25 |
| factera | 1 | 6.8 | 0.125 | 35 |
| seeksv | 3 | 69.7 | 0.375 | 973 |
| svaba | 7 | 11.7 | 0.875 | 73 |
| breakdancer | 6* | 1.7 | 0.75 | 2011 |
| breakmer | 7 | 123.6 | 0.875 | 121 |

### Public data sets

To further verify the feasibility of FirstSV in general, public data set from NCBI, including 110 samples with known structural variations, was downloaded. Details about the sample information were described in citation^18^, and the structural variations in samples were all validated by experiments. All samples were fragmented using sonication to mean insert size of 250bp and were purified using Agencourt AMPure XP beads. The 100bp, paired-end sequences were generated using a HiSeq 2500 platform in rapid run mode. All the sequencing data analyzed in this study have been submitted to the NCBI Short Read Archive databank (SRA, http://www.ncbi.nlm.nih.gov/sra) under accession number SRP042598 (SRA).

After being downloaded, raw sequences were preprocessed, followed with an above-mentioned standard process to get the BAM files. Then, 110 samples were analyzed using FirstSV and other software pieces including seeksv, svaba, factera, breakmer, breakdancer, meerkat, and crest. The detection rate of FirstSV was 93.6%, seeksv 45.5%, svaba 79.1%, and factera 64.5%. As shown in Table 3, eight variations were not detected by FirstSV, in which one was due to incomplete raw data downloaded from the public data set; and the other seven false negatives were due to insufficient sequencing depth in the region of breakpoints. In other words, the number of support reads did not meet FirstSV default threshold (the default was 5 reads). When this threshold is lowered, FirstSV could detect all of them.

**Table 3.**
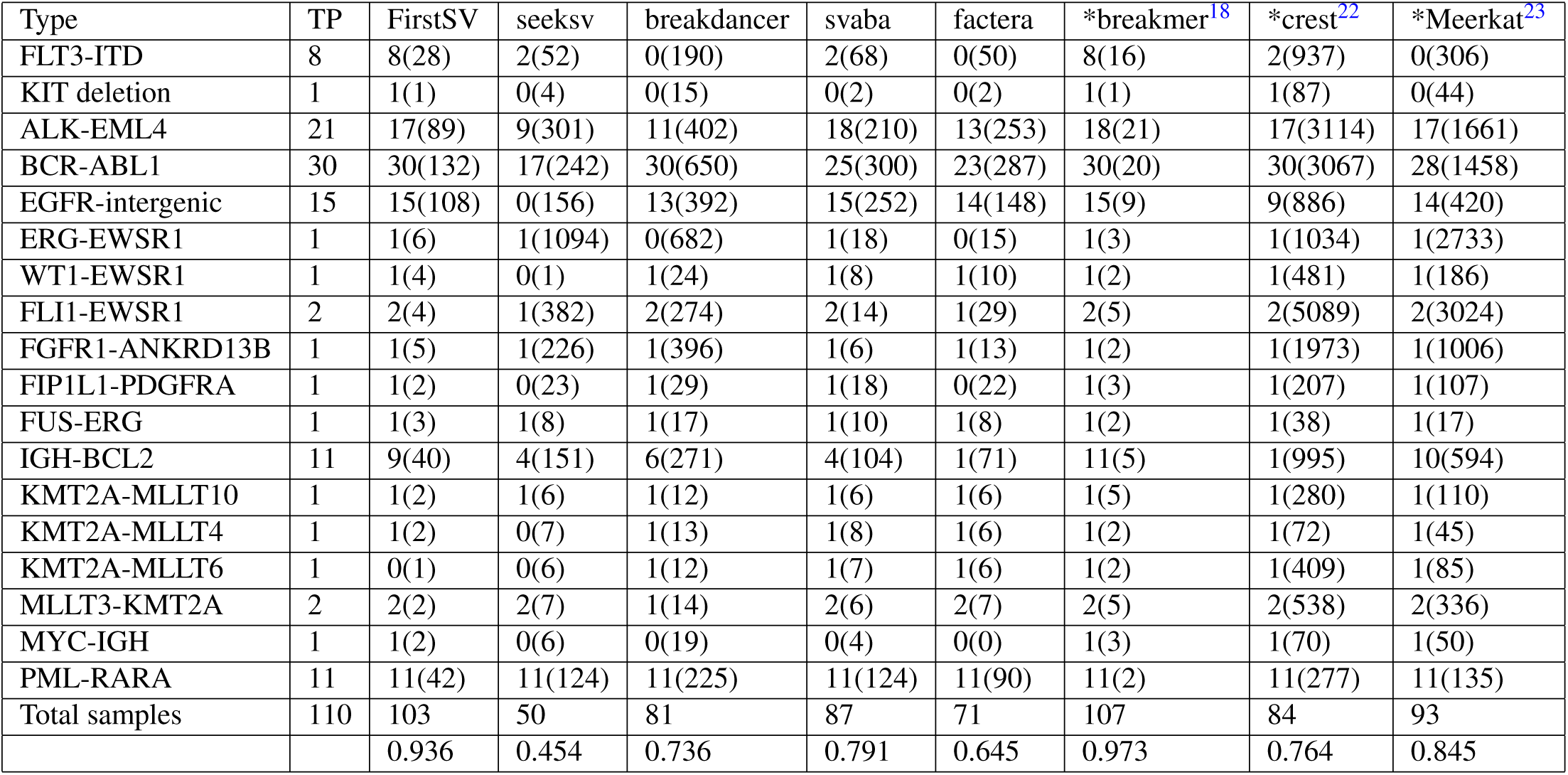
Detections of SVs in public data. 2nd column: True positive (TP for short) represented the number of samples with true structural variations. The figures in brackets represented the total number of additional calls, which are false positives.

**Table 4.** SV detections of simulated data. * represented simulated data with different coverages (columns 2-8). The other columns (9-15)represented simulated data with different mutation frequency (the coverage was about 5000). The figures represented the number of SV detected. The true positive figures for deletion, inversion, and translocation is 34, 38, and 75 correspondingly.

| Software | Sequencing Coverage |  |  |  |  |  |  | Frequency of Known Structural Variations |  |  |  |  |  |  |
| --- | --- | --- | --- | --- | --- | --- | --- | --- | --- | --- | --- | --- | --- | --- |
|  | 10X* | 20X* | 50X* | 100X* | 500X* | 1000X* | 5000X* | 0.001 | 0.005 | 0.01 | 0.05 | 0.2 | 0.5 | 1 |
| deletions |  |  |  |  |  |  |  |  |  |  |  |  |  |  |
| FirstSV | 34 | 34 | 34 | 34 | 34 | 34 | 34 | 34 | 34 | 34 | 34 | 34 | 34 | 34 |
| svaba | 10 | 10 | 10 | 9 | 6 | 2 | 2 | 5 | 4 | 9 | 9 | 8 | 8 | 10 |
| breakdancer | 30 | 33 | 34 | 34 | 34 | 34 | 34 | 32 | 32 | 32 | 32 | 32 | 32 | 32 |
| breakmer | 27 | 27 | 27 | 27 | 27 | 27 | 27 | 27 | 27 | 27 | 27 | 27 | 27 | 27 |
| inversions |  |  |  |  |  |  |  |  |  |  |  |  |  |  |
| FirstSV | 38 | 38 | 38 | 38 | 38 | 38 | 38 | 38 | 38 | 38 | 38 | 38 | 38 | 38 |
| svaba | 14 | 8 | 6 | 8 | 5 | 4 | 8 | 9 | 4 | 9 | 10 | 11 | 9 | 8 |
| breakdancer | 34 | 37 | 37 | 38 | 38 | 38 | 38 | 35 | 35 | 35 | 35 | 35 | 35 | 35 |
| breakmer | 29 | 29 | 29 | 29 | 29 | 29 | 29 | 29 | 29 | 29 | 29 | 29 | 29 | 29 |
| translocation |  |  |  |  |  |  |  |  |  |  |  |  |  |  |
| FirstSV | 75 | 75 | 75 | 75 | 75 | 75 | 75 | 75 | 75 | 75 | 75 | 75 | 75 | 75 |
| svaba | 29 | 17 | 20 | 20 | 14 | 15 | 17 | 12 | 10 | 15 | 22 | 21 | 18 | 17 |
| breakdancer | 69 | 69 | 69 | 69 | 69 | 65 | 54 | 63 | 63 | 63 | 63 | 65 | 65 | 64 |
| breakmer | 62 | 64 | 65 | 65 | 65 | 65 | 65 | 62 | 62 | 64 | 64 | 65 | 65 | 65 |
| Additional calls |  |  |  |  |  |  |  |  |  |  |  |  |  |  |
| FirstSV | 0 | 0 | 0 | 0 | 5 | 11 | 80 | 0 | 0 | 0 | 0 | 2 | 2 | 2 |
| svaba | 0 | 1 | 2 | 1 | 0 | 0 | 1 | 1 | 0 | 0 | 1 | 0 | 0 | 1 |
| breakdancer | 36 | 47 | 43 | 53 | 45 | 116 | 57 | 10 | 8 | 9 | 11 | 17 | 15 | 16 |
| breakmer | 4 | 2 | 7 | 6 | 14 | 23 | 67 | 4 | 1 | 2 | 5 | 7 | 6 | 1 |

From Table 3, one can see that FirstSV still has a small number of false negative cases (a total of 7 out of 110 cases, 4 in ALK-EML4 fusion, 2 in IGH-BCL2 fusion, and 1 in KMT2A-MLLT6 fusion). Actually, all of these mutations can be detected by FirstSV through threshold adjustments, but this would introduce more false positives. The default parameters we used here actually means a trade-off between lowering the false-positive rate and decreasing the false-negative rate the same time.

## Discussion

The low specificity of previous SV detection methods highlighted the need for more accurate structural variations detection approaches. Our original intention was to design a software for SV detection of cfDNA data. We first used the softclip reads analysis as the length of the cfDNA fragment was close to the sequence read length. The depth of cfDNA sequencing was much higher than that of WGS or WES sequencing and that deems most of the existing SV detection methods inappropriate. Specifically, performing initial filtering of candidate areas is necessary to reduce the computational burden. Just analyzing softclip reads alone, however, was not sufficient when the insert size was much longer than read length. Thus, we took into account discordant reads signature to solve this problem. The final resulted hybrid program, FirstSV, was proven to be a higher sensitive and specific method for the detection of breakpoints in targeted sequencing data.

Moreover, FirstSV could be applied to any other sequencing data sources including paired-end data, and single-end data. Although originally implemented for structural variations detection in cfDNA applications, FirstSV can also be applied toward broader applications. For example, FirstSV can be used to analyze structural variations in patient tissue samples, and that have been proven through our many validations.

## Methods

The input of FirstSV is a BAM file. FirstSV could handle pair-end data or single-end data. When dealing with pair-end data, both of the softclip reads signature and the discordant reads signature were considered. FirstSV first extracted discordant reads and softclip reads to construct abnormal sequence datasets. Next, potential breakpoints were confirmed based on the softclip reads signature and the discordant reads signature. Then the sequences near the potential breakpoints were assembled to contigs and the contigs were aligned to the reference genome to obtain the breakpoints. Flowchart of our data processing was shown in Figure 2.

**Figure 2.**
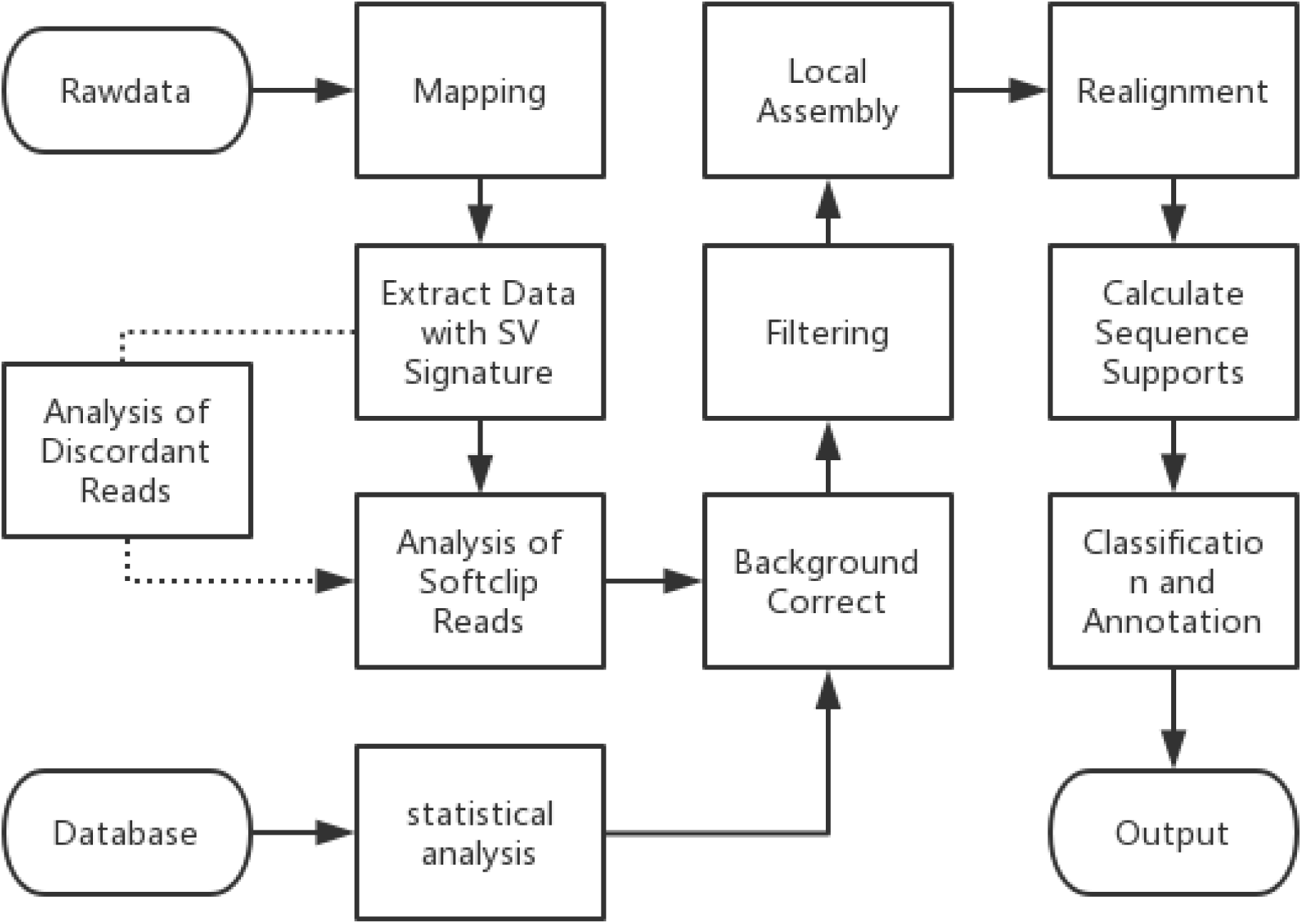
Schematic of FirstSV.

### Data preprocessing

Data was aligned to the hg19 reference genome using the mem algorithm of BWA(0.7.13)^19^, and the BAM files were sorted using samtools (1.3)^20^. Duplicate reads were identified and removed using Picard tools. The alignments were further refined using the GATK tool for localized realignment around indel sites. Recalibration of the quality scores was performed using GATK tools. The resulted BAM files could be used for subsequent structural variations detection. The above tools were ran by default parameters.

### Softclipreads extraction

FirstSV extracted softclip reads of which the alignment quality was higher than Q20 from the BAM file. The fasta format dataset was constructed for subsequent analysis using the extracted sequences. Sequences with softclipped length above a certain threshold is needed in order to map them to the reference genome.

### Discordant reads analysis

FirstSV also uses the discordant reads signature when dealing with paired-end data, and extracted the information of all discordant reads from the BAM file. Potential breakpoints were confirmed by clustering the information of each pair of discordant reads. We hypothesize that if two pairs of discordant reads were located near the same potential breakpoints, the potential breakpoints might be originated from the same structural variation. Based on this hypothesis, FirstSV clustered all the inconsistent sequences, in which each of clusters contained the location information of a region on the genome and the number of the inconsistent sequences contained in the region.

The structural variation detection algorithms were different for different sequencing data types. Discordant read signature is especially useful for pair-end reads. As shown in Figure 2, to make our method suitable for different data types, the analysis of discordant reads signature was added later into FirstSV. Softclip reads analysis was the main algorithm.

### Softclip reads analysis

Discordant reads analysis usually fails to get the exact potential breakpoint location and just give a potential region where the potential breakpoint might existed. Discordant reads analysis was applied to get the approximate location of the region to extract softclip reads for detection efficiency. However, risk of missing detection could not be avoided. To solve this problem, FirstSV analyzed all valid softclip reads in the BAM file. The valid softclip reads meant that the softclipped part of the sequence can be realigned to the reference genome and the alignment quality was higher than Q20. If the softclipped part could not realign to reference genome uniquely, the softclip reads were not considered valid. As mentioned above, the location of potential breakpoints was confirmed by extracting information of softclip reads alignment. The potential breakpoints were filtered by softclip reads supports and discordant reads supports with a default threshold of 5. In addition to these basic filters, additional filters, such as GC content filters and sequence complexity filters, were introduced.

### GC content filtering

We assume that the GC content of sequences near breakpoints of the real structural variations is within a certain range so that a portion of the false positives can be filtered out with the GC content threshold filter. Based on this assumption, we analyze the GC content of sequences near the breakpoint of structural variations in COSMIC and sequences in the regions that often contain pseudo structural variations. The version of COSMIC was v79^21^.

The average GC content of all the sequences near breakpoints in COSMIC was 0.4 (Figure 3). By analyzing the data of 35 healthy human samples, the GC content of the sequence near the potential breakpoint was calculated. We chose 100 values ranging from 0.01 to 1 as our potential threshold to filter the data of 35 healthy samples and COSMIC sequence. Then false positive rates and true positive rates for each threshold were calculated to draw a ROC curve. On the ROC curve, the point closest to the top left of the coordinate diagram is the critical value with high sensitivity and specificity. The threshold of GC content filter was determined ranging from 0.27 to 0.73. In other words, the detected SVs in this GC interval would be retained. In order to discuss whether the number of samples would affect the determination of the threshold, we selected 1, 2, 3, 4, 5, 10, 15, 20, 30, 35 samples to determine the threshold value with the same method. After analysis, the thresholds determined by methods using more than 5 samples were the same. So at least 5 normal samples were needed to determine an ideal threshold.

**Figure 3.**
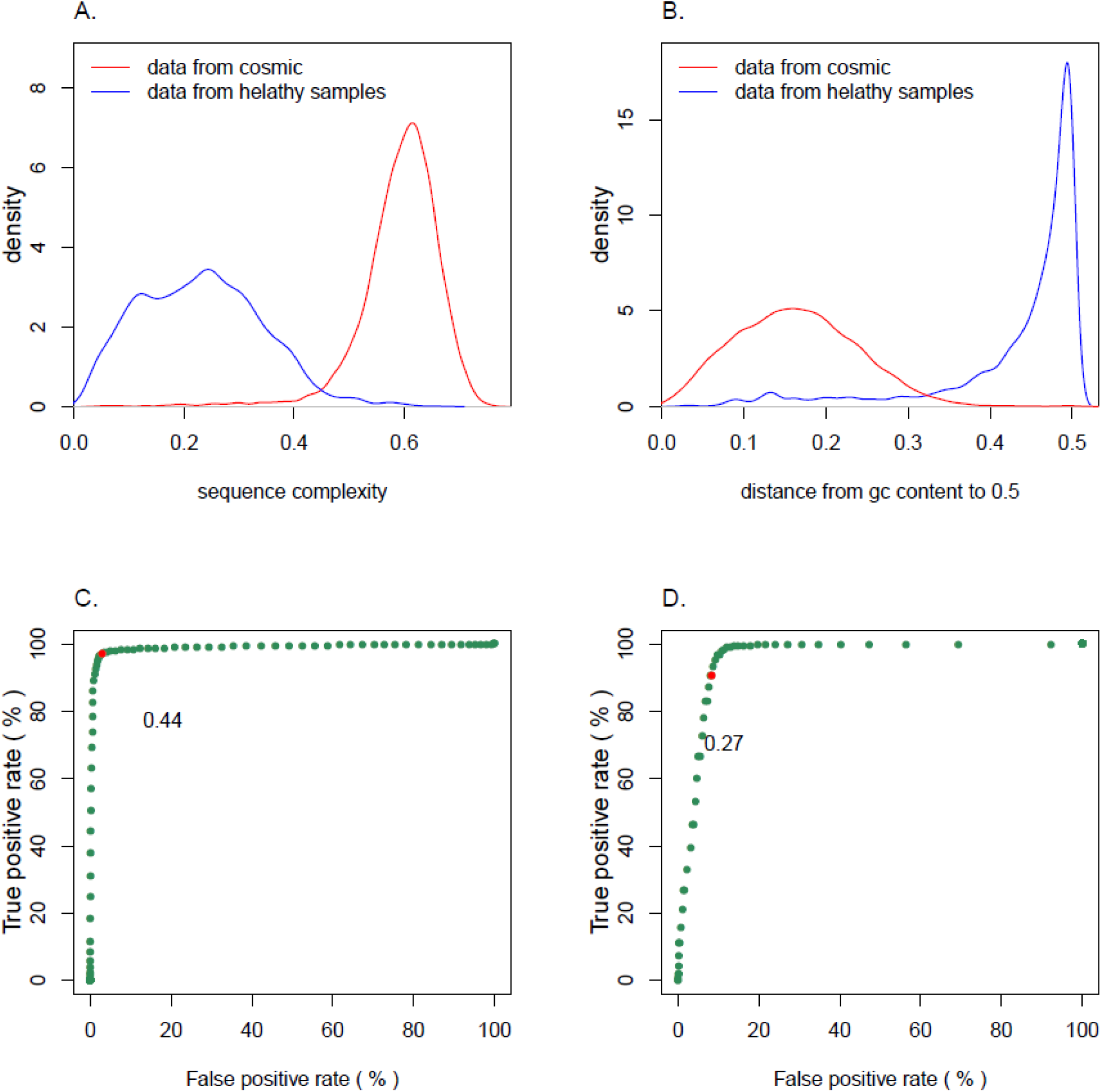
Determination of the threshold. To determine the threshold for filtration, 35 healthy samples were selected for analysis. The optimal threshold was calculated by the ROC curve. (A) showed the distribution of the sequence complexity. The vertical axis was the density and the horizontal axis was sequence complexity. (B) showed the distribution of the distance from GC content to 0.5. The vertical axis was the density and the horizontal axis was the distance from GC content to 0.5. (C) showed the ROC curve of sequence complexity. The threshold of sequence complexity is 0.44. (D) showed the ROC curve of GC content. The threshold of GC content was ranged from 0.27 to 0.73.

### Sequence complexity filtering

Sequences with low complexity often introduce errors when they were mapped to the reference genome, and this type of mapping error would interfere with the detection of SVs. To avoid these false positives caused by mapping errors, sequence complexity filtering was applied in FirstSV. There was a large difference in the complexity of the sequence on either side of the breakpoint location at the fake breakpoint locations. The complexity threshold was determined by the COSMIC database data and potential breakpoint data from 35 healthy individuals. The method to determine the complexity threshold was the same as GC content filtering.

### Background correction

Background errors from target capture panel affected the detection of structural variations. 35 cfDNA samples from healthy people were chosen to correct background errors.

The locations of potential breakpoints were obtained through the previous steps, and then analyzed to find areas on the test panel in which the number of potential breakpoints were more than others. The background distribution of variation frequency was then estimated based on the potential breakpoints detected in these areas. To increase the detection threshold of error-prone areas on the panel and avoid most of the false positives in these areas, the background distribution was used to correct the supports of the potential breakpoints.

### Local assembly

Sequences based on the filtered breakpoints were extracted and assembled. The local assembly algorithm in FirstSV was based on the Fermi package [reference here], which was an overlap layout consensus algorithm. In the package, BFC algorithm was used to correct sequence errors before assembly. FM-index algorithm, suitable for assembling small data sets with high efficiency and keeping polyploids with parameter adjustments, was used to establish the overlay graph.

A piece of local assembly algorithm was written in C, and it handled BAM files and breakpoints files directly. Based on the location of breakpoints, it extracted the nearby inconsistent sequences and mismatched sequences directly from the BAM file. The default range was 150bp (as the sequencing length was 150bp). If the sequences extracted from potential breakpoints were not enough to assemble, the local assembly algorithm simply gave up the assembly. Moreover, the assembly algorithm removed false positives caused by mapping errors and corrected breakpoint locations. The contigs were stored in the file of fastq format.

### SV verification

BWA tool was applied to align contigs to the reference genome to determine breakpoints one more time in FirstSV. Breakpoints included in a list of potential breakpoints would be considered as the final breakpoints. Then, the BLAT tool was used to search the inconsistent sequences dataset established in the first step for the support of pair of breakpoints. The sequence would be considered a support if the sequence contains the breakpoint position and the alignment is above 90% length of the contig.

### Data Availability

The datasets generated during and/or analysed during the current study are available from the corresponding author on reasonable request.

## Acknowledgements

We thank LiHeng for contributions to an version of the Fermi algorithm used in the FirstSV source code. The results published here are in part based upon data generated by Dana-Farber Cancer Institute.

## Author contributions statement

J.S. and Q.Y.Z. wrote FirstSV code, conducted bioinformatics analysis, and wrote the manuscript. R.L.W. helped the article in a biological perspective and wrote the manuscript. X.L. sorted data and draw figures. Y.H.T provided guidance for the manuscript. All authors reviewed the manuscript.

## Additional information

### Publisher’s note

Springer Nature remains neutral with regard to jurisdictional claims in published maps and institutional afliations

